# Representing context and priority in working memory

**DOI:** 10.1101/2023.10.24.563608

**Authors:** Quan Wan, Adel Ardalan, Jacqueline M. Fulvio, Bradley R. Postle

## Abstract

The ability to prioritize among contents in working memory (WM) is critical for successful control of thought and behavior. Recent work has demonstrated that prioritization in WM can be implemented by representing different states of priority in different representational formats. Here, we explored the mechanisms underlying WM prioritization by simulating the double serial retrocuing (DSR) task with recurrent neural networks (RNNs). Visualization of stimulus representational dynamics using principal component analysis (PCA) revealed that the network represented trial context (order of presentation) and priority via different mechanisms. Ordinal context, a stable property lasting the duration of the trial, was accomplished by *segregating* representations into orthogonal subspaces. Priority, which changed multiple times during a trial, was accomplished by *separating* representations into different manifolds within each subspace. We assessed the generality of these mechanism by applying dimensionality reduction and multiclass decoding to fMRI and EEG datasets and found that priority and context are represented differently along the dorsal visual stream, and that behavioral performance is sensitive to trial-by-trial efficacy of priority coding, but not context coding.

---

One of the hallmarks of working memory (WM) is its ability to flexibly prioritize among its contents in the service of the current behavioral goal. For example, say that you’ve just completed a talk at a conference, and you see two people simultaneously approaching each of two microphones to ask a question. You turn to the moderator and wait for them to indicate who will ask the first question, and based on this your shift of gaze is guided by your memory of the location of the cued microphone. To study prioritization in WM, one line of work has made extensive use of the double serial retrocuing (DSR) task, in which two sample items are initially presented and “remembered”, followed by a blank “no-action” delay, then a retrocue indicating which of the two memorized items will be tested by an impending memory probe (see Figure 1A for an example). This item is said to take on the status of prioritized memory item (PMI). Because the item that was not cued may be tested later in the trial, however, it cannot be dropped from memory (i.e., “forgotten”), so it takes on the status of unprioritized memory item (UMI) until the PMI is tested. Subsequently, a second retrocue indicates, unpredictably, which item will be tested by a second memory probe; thus, either item can take on the status of PMI during the second half of the trial. An initial set of studies applying multivariate pattern analysis (MVPA) decoding to fMRI and EEG data from subjects performing the DSR task failed to find evidence for an active representation of the UMI, giving rise to the idea that it might be held in an “activity-silent” state (Larocque et al., 2014; LaRocque et al., 2017; Lewis-Peacock et al., 2011; Rose et al., 2016). More recently, however, studies using variants of the DSR task (with fMRI; van Loon et al., 2018; Yu, Teng & Postle, 2020) and the 2-back WM task (with EEG; Wan et al., 2020) have provided evidence for an active trace of the UMI that undergoes a transformation relative to the representational format of the PMI. Specifically, the UMI can produce significantly below-baseline MVPA decoding (van Loon et al., 2018) and “opposite” reconstruction with multivariate inverted encoding modeling (IEM; Wan et al., 2020; Yu, Teng & Postle, 2020).

**Figure 1.**
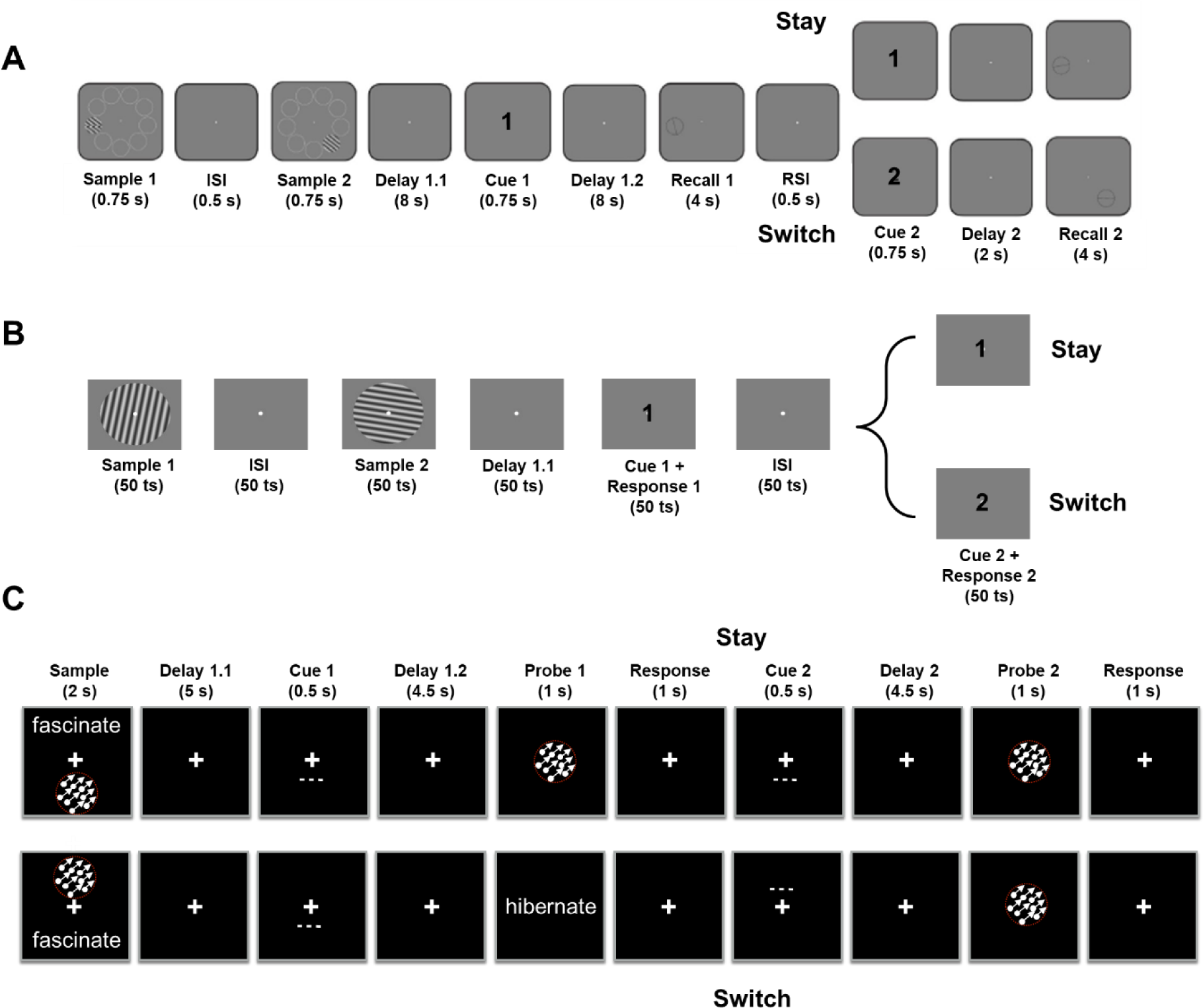
Experimental procedure for (A) The fMRI task, (B) the RNN task and (C) the EEG task. Figures adapted, with permission, from Yu, Teng and Postle (2020; panel A), Fulvio and Postle (2020; panel C).

As an initial step toward better understanding the priority-based representational transformations observed in neuroimaging data (van Loon et al., 2018; Wan et al., 2020; Yu, Teng & Postle, 2020), we had trained recurrent neural networks (RNN) with a long short-term memory (LSTM) architecture to perform the 2-back WM task (Wan et al., 2022). Visualization of LSTM hidden layer activity using principal component analysis (PCA) had confirmed that stimulus representations in RNNs also undergo representational transformations when transitioning between priority states. Specifically, demixed (d)PCA of these data had identified two representational trajectories, one within a UMI-specific subspace and the other a PMI-specific subspace, both undergoing a reversal of stimulus coding axes. Having thus observed similar priority-based transformational dynamics in the human brain and in RNNs, we speculated that this type of transformation might be a computationally rational way to meet the competing demands of retaining information in WM while simultaneously preventing it from interfering with concurrent behavior (Wan et al., 2022).

Whereas in Wan et al. (2022) we simulated the 2-back task, the results presented here were prompted by results of RNN simulation of the DSR. This was important to do because although the n-back task has been important for the study of many aspects of WM, it is poorly suited for the study of the flexible control of behavior with WM. This is because the n-back is a continuous performance task in which each item follows the same functional trajectory. For the 2-back, for example, each item *n* first serves as a memory probe against which to compare one’s memory for item *n - 2*, then transitions to UMI (while *n + 1* is compared with the memory of *n – 1*), then transitions to PMI (for its comparison with item *n + 2*), then becomes no-longer-relevant and can be dropped from WM. The DSR, in contrast, does require online, flexible control, because the identity of the two retrocues can’t be predicted prior to their onset. Unexpectedly, however, it is a different property of the DSR task that motivated the work presented here.

At the beginning of each trial of DSR, sample items can either be presented simultaneously or serially. When items are presented simultaneously, they necessarily each appear at a different location, and it is an item’s unique location that is used by the retrocue to designate it the PMI. Thus, the location at which an item appears serves as critical trial-specific context. When items are presented serially, they can appear at the same or different locations, and when they appear at the same location (as they did in Yu, Teng & Postle, 2020) the retrocue must designate the prioritized item by referring to the order in which it was presented (i.e., “first” or “second;” the item’s ordinal context). When we simulated the DSR from Yu, Teng and Postle (2020), the unexpected finding was that the representation of the first sample item underwent a dramatic transformation upon the onset of the second item (i.e., prior to the designation of priority, which would be indicated by the retrocue). Specifically, whereas it had been represented in a subspace defined by the first two principal components of a PCA applied to the hidden layer or the RNN, it was displaced from this subspace by the representation of the second item, and shunted to a new subspace defined by the third and fourth principal components of the PCA. This finding caused us to reconsider our interpretation of the transformational dynamics observed in the 2-back task (Wan et al., 2022), because an item’s functional trajectory during that task confounds priority with context. That is, while an item has the status of UMI it also has the contextual status of item-that-was-presented-most-recently (i.e., “1-back”), and when it then transitions to the status of PMI its context simultaneously transitions to item-that-was-presented-2-back. The aims of this report, therefore, are two-fold. One is to explore, at the computational level, how context-based representational transformations may differ from priority-based transformations. This will be carried out via RNN simulations. The second is to assess how these two properties, context and priority, might differ in the way they influence behavior, and in the way they are represented in the brain.

## Methods

The data presented here derive from three sources: RNN simulations of a double serial retrocuing (DSR) task; reanalysis of data from an EEG study of DSR; and reanalysis of an fMRI study of DSR.

### Participants

#### EEG

The EEG data set is from 12 healthy young adults (5 females, average age = 21.7 ± 3.2 years, all right-handed), as described in detail in Fulvio and Postle (2020). This *N* was double that of a previous EEG study for which MVPA decoding results yielded informative prioritization effects (Rose et al., 2016), and so was deemed satisfactory for the analyses to be carried out here.

#### fMRI

The fMRI data set is from 13 healthy young subjects (10 females, average age = 21.1 ± 4.5 years, all right-handed), as described in detail by Yu, Teng, and Postle (2020). Because IEM analyses of these fMRI data had yielded informative prioritization effects, this *N* was deemed satisfactory for the analyses to be carried out here.

### Behavioral tasks

#### Recurrent Neural Network (RNN) models

The training task was modeled after the fMRI task (Yu, Teng & Postle, 2020; Figure 1B). Stimuli were randomly drawn from a pool of oriented gratings that covered the continuous range from [0°, 180°) interval (*Sample 1*: φ and *Sample 2*: θ). Stimulus location was not simulated, and it was possible for φ and θ to take on the same orientation. Each trial began with the presentation of *Sample 1* (50 timesteps) followed by an interstimulus interval (ISI, i.e., blank delay; 50 timesteps) followed by *Sample 2* (50 timesteps) followed by *Delay 1.1* (50 timesteps) followed by *Cue 1/Response 1* (50 timesteps; the response window was the duration of *Cue 1*). Next came another ISI (50 timesteps) followed by *Cue 2/Response 2* (50 timesteps). *Cue 2* matched (“stay”) or did not match (“switch”) *Cue 1*, unpredictably, and equal number of times.

#### fMRI: DSR with ordinal and location context, and recall probes

Stimuli were drawn from a pool of 9 oriented gratings that evenly covered the range from 0° to 179°, and could be presented at one of 9 locations that, each at a distance of 8° of visual angle from central fixation, evenly covered the range of possible locations from 0° to 359° of polar angle. Each trial began with the presentation *Sample 1* (.75 sec) followed by an ISI (.5 sec), followed by *Sample 2* (.75 sec), followed by *Delay 1.1* (8 sec), followed by a centrally presented digit (“1” or “2,” *Cue 1*; .75 sec). After the ensuing *Delay 1.2* (8 sec), a recall dial appeared at the location that had been occupied by the PMI, and the subject had 4 sec to rotate it to match their memory of that item’s orientation. Subsequently, after a brief unfilled interval (.5 sec), a second centrally presented digit (“1” or “2,” *Cue 1*; .75 sec) indicated the item to be tested, after *Delay 2* (2 sec), at *Recall 2* (4 sec). *Cue 2* matched (“stay”) or did not match (“switch”) *Cue 1*, unpredictably, an equal number of times (Figure 1A).

Because the location of the recall dial indicated the item to be recalled, a possible strategy would be to ignore the cues and simply behave based on the location of the recall dial. However, this strategy was discouraged due to an important detail of the procedure. On each trial, the orientation and the location of each stimulus were selected at random (with replacement), and independently. Thus, on each trial there was a *p* = .11 chance that the second sample would have the same orientation as the first and, independently, a *p* = .11 chance that the second sample would appear at the same location as had the first. These contingencies encouraged subjects to not wait for the onset of the recall dial to recall the orientation of the PMI and, indeed, patterns of priority-related transformation of the UMI during *Delay 1.2*, as assessed by IEM, confirmed that subjects used the ordinal cue to guide their behavior (Yu, Teng, and Postle, 2020).

#### EEG: DSR with location context and recognition probes

Each trial began with the simultaneous presentation of two sample items, one drawn from each of two out of three possible categories (faces, direction of dot motion, and words), one appearing above and one below central fixation (2 sec; Figure 1C). The samples were replaced by a central fixation symbol (“+”) during an initial delay (*Delay 1.1*; 5 sec), followed by a dashed line appearing at one of the two sample locations (.5 sec), indicating that that item would be the first to be tested (*Cue 1*). After a second delay (*Delay 1.2;* 4.5 sec), during which the cued item had the status of prioritized memory item (PMI) and the uncued item the status of unprioritized memory item (UMI), an image serving as a recognition probe appeared centrally, and was either identical to the PMI (“match,” *p* = .5), drawn from the same category but a different exemplar than the PMI (“nonmatch,” *p* = .3), or identical to the UMI (also “nonmatch”, *p* = .2; *Probe 1;* 1 sec). *Probe 1* was replaced by the fixation symbol (*Response 1*; 1 sec), and a response was required during the 2 sec spanning Probe 1 and Response 1. Next a dashed line appeared at one of the two sample locations (*Cue 2*; .5 sec), thereby designating the PMI for the following *Delay 2* (4.5 sec), then *Probe 2* (1 sec) then *Response 2* (*1 sec*). ITI varied from 2-4 sec.

Data were collected during three sessions, each on a separate day, with each session comprising eight 30-trial blocks, alternating between blocks of DSR and a single retrocue task (results from single retrocue task not presented here). During each block *Cue 1* appeared unpredictably at the “up” or “down” location an equal number of times and, orthogonal to *Cue 1* location, *Cue 2* appeared, unpredictably, at the same (“Stay,” Figure 1C, top row) or opposite (“Switch,” Figure 1C, bottom row) location as had *Cue 1* an equal number of times. Balanced across cue conditions, spTMS was delivered 2-3 sec after the offset of *Cue 1* on 50% of trials and, orthogonally, after the offset of *Cue 2* on 50% of trials. Note that the EEG data used for the “transformation efficacy analyses” (see the “Analysis procedures” section below) include both epochs with and without spTMS.

### Experimental procedures

#### RNN

Stimulus orientations were fed into the network via 32 orientation-tuned input units whose preferred orientations spanned the full 180° range, and whose response properties were based on V1 orientation-selective neurons (Teich & Qian, 2003; Figure 2). A 33^rd^ input unit was used for retrocue, inputting “0” on each non-cue timestep and a “1” or “-1” (indicating “1st” or “2nd,” respectively) during each cue timestep. The two output units were trained to produce cos(2*x*) and sin(2*x*) (where *x* was either θ or φ depending on the cue) so that the 0° orientation had the same output as the 180° orientation (Figure 2).

**Figure 2.**
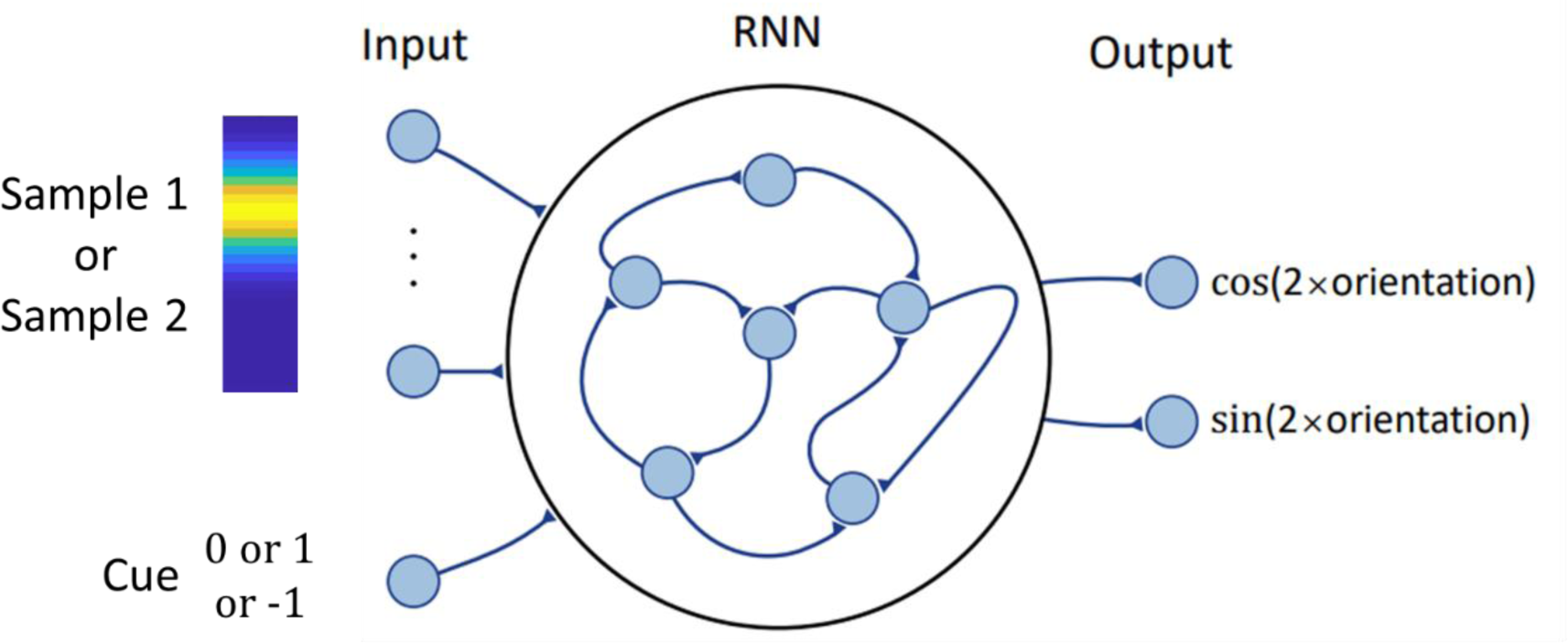
RNN input and architecture. Top left illustrates input of a stimulus with an angular value corresponding to the peak magnitude of this 32-dimensional vector; bottom left illustrates that at each timestep the value of the input to the cue input unit was *0*, *1*, or *-1*.

Our network had 100 fully-connected recurrent units and the dynamics *u*_*i*_(*t*) of each recurrent unit were governed by the following standard continuous-time RNN equations:

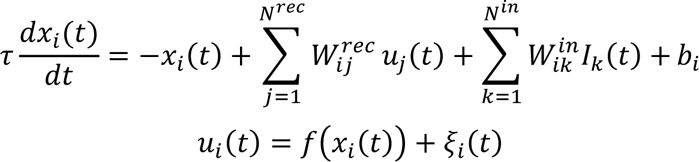

for *i* = 1, …, *N*^*rec*^. We introduced nonlinearity using the rectified linear unit (ReLU) function *f*(*x*) = max(0, *x*). Each recurrent unit received input from other units via recurrent connections with weights specified by the matrix *W*^*rec*^, initialized orthogonally (Saxe et al., 2013). In addition, these units received external input *I*(*t*) to the RNN via weights specified by the matrix *W*^*in*^. Each unit carried two sources of bias: (1) *b*_*i*_, learned during training, and (2) *ξ*_*i*_(*t*), which represented intrinsic noise in the network and took the form of white Gaussian (sampled independently at each timestep) with zero mean. We simulated the approximate network dynamics using the Euler method for *T* = 350 timesteps, each having a duration *τ*/10 (Mante et al., 2013). We chose *dt*/*τ* = 0.1 similar to (Cueva et al., 2021); e.g., *dt* = 10 ms and *τ* = 100 ms, which would make the time scale of our simulations close to that of the fMRI experiment. The outputs *y*_j_(*t*) were then generated by combining the activities of the recurrent units based on:

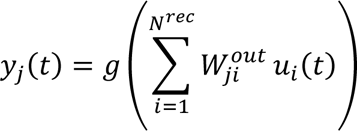

where *g* is the tanh activation function.

We optimized the network parameters *W*^*in*^, *W*^*rec*^, *b* and *W*^*out*^ to minimize the mean squared error between the target outputs and the network outputs:

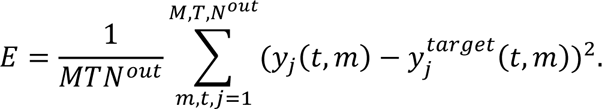

Parameters were updated with the Adam stochastic gradient descent (SGD) algorithm (Kingma & Ba, 2014) and each network was trained for 10,000 epochs.

#### fMRI

For each subject regions of interest (ROIs) were defined, both anatomically and functionally, for eight regions: early visual cortex (EVC, V1 and V2 merged), IPS0-through-IPS5 (6 ROIs), and FEF (all ROIs cover both hemispheres). First, anatomical ROIs were defined by extracting masks from the probabilistic atlas of Wang and colleagues (2015) and warping them to each subject’s structural scan in native space. To identify task-related activity, we modeled each epoch of the task with 6 boxcar regressors in a general linear model (GLM) – *Sample* (2 sec), *Delay 1.1* (8 sec), *Delay 1.2* (8 sec), *Recall 1* (4 sec), *Delay 2* (2 sec), and *Recall 2* (4 sec) convolved with a canonical hemodynamic response function and we also included covariates to control for motion. We proceeded to create anatomically constrained functional ROI for bilateral EVC by selecting the 500 voxels inside the V1-2 anatomical ROI with the strongest loading on the *Sample* regressor and for bilateral IPS0-5 and FEF by separately selecting the 500 voxels inside each of IPS0-5 and FEF anatomical ROIs, with highest loading on the *Delay 1.2* regressor.

#### EEG data collection

The experimental procedure from the experiment reported by Fulvio and Postle (2020) entailed recording the EEG with concurrent delivery of single pulses of transcranial magnetic stimulation (spTMS) on half of the delay periods of the DSR task. However, because the original report only included behavioral results (with and without spTMS), here we detail the EEG methods.

EEG was recorded with a 60-channel cap and TMS-compatible amplifier, equipped with a sample-and-hold circuit that held amplifier output constant from 100 μs before stimulation to 2 ms after stimulation (NexStim eXimia, Helsinki, Finland). Electrode impedance was kept below 5 kΩ. The reference electrode was placed superior to the supraorbital ridge. Eye movements were recorded with two additional electrodes, one placed near the outer canthus of the right eye, and one underneath the right eye. The EEG was recorded between 0.1 and 350 Hz at a sampling rate of 1450 Hz with 16-bit resolution.

Data were processed offline using EEGLAB (Delorme & Makeig, 2004) with the TMS-EEG signal analyzer (TESA) open-source EEGLAB extension (Mutanen et al., 2020; Rogasch et al., 2017) and Fieldtrip (Oostenveld et al., 2010) toolboxes in MATLAB. The pipeline followed the TMS-EEG analysis pipeline (http://nigelrogasch.github.io/TESA/). Then, electrodes exhibiting excessive noise were removed and the data were epoched to -12 s to 8 s around the first spTMS event tag (*Delay 1.2*) and -4.5 s to 4.5 s around the second spTMS event tag (*Delay 2*). The data were downsampled to 500 Hz. In order to minimize the TMS artifact in the EEG signal, the data were interpolated using a cubic function from -2 to 30 ms around the TMS pulse, and this interpolation was also carried out on delay periods on which TMS was not delivered. (For delay periods for which no spTMS was delivered (“spTMS-absent”), a dummy spTMS event tag was added at a latency that matched the most recent spTMS-present delay period.) The data were bandpass filtered between 1 and 100 Hz with a notch filter centered at 60 Hz. Independent component analysis (ICA) was used to identify and remove components reflecting residual muscle activity, eye movements, blink-related activity, residual electrode artifacts, and residual TMS-related artifacts. A spherical spline interpolation was applied to electrodes exhibiting excessive noise. Finally, the data were re-referenced to the average of all electrodes that were included in the ICA.

The present analyses included EEG data from all delay periods (i.e., averaging data from spTMS-present and spTMS-absent trials and ignoring this factor).

### Analysis procedures

#### PCA visualization of the RNN hidden layer activity

We extracted from each network the activity of the 100 recurrent units from all 1000 testing trials and used PCA to project these 100-dimensional activity patterns onto the four dimensions accounting for the most variance across all training trials separately for each timestep. We then visualized the representations of each *Sample 1* and *Sample 2* by plotting the dimensionality-reduced activity across the 350-timestep time course of a trial, and coloring the activity patterns according to stimulus identity, separately, in three 2D plots (PC1-2, PC2-3 and PC3-4).

In addition, we plotted the effective dimensionality (ED) of the data at each timepoint, which is the equivalent number of orthogonal dimensions that would produce the same overall pattern of covariation (Del Giudice, 2021). It is calculated using the following formula:

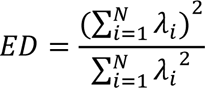

where λ_*i*_s are the eigenvalues of the covariance matrix of the *N* recurrent units’ activities at a certain time point.

Transformational efficacy analyses on EEG and fMRI data

The PCA visualizations of RNN activity revealed representational dynamics, such that stimulus information was represented differently as a function of context (1^st^ or 2^nd^) and as a function of cue identity (essentially priority (PMI or UMI/IMI) for each stimulus). To assess the functional relevance of these two coding schemes for human behavior, we assessed trial-by-trial variation in the efficacy of context-based and priority-based transformations, and determined for each whether this variability related to trial-by-trial variation in behavior.

For the representation of context, we first calculated a template stimulus representational format for each subject by averaging the neural activity for each context status (“1^st^” or “2^nd^” for fMRI; “up” or “down” for EEG) over a time window corresponding to *Delay 1.1*, across all trials. (For the remainder of this section, for simplicity, we will only refer to ordinal context.) To these two windowed averages we applied demixed principal component analysis (dPCA; refer to Wan et al. 2022 for methodological details) to derive the first two demixed principal components (PCs), thereby constructing a *Sample 1* template subspace and a *Sample 2* template subspace. We then projected individual trial activity from the same time window into the template subspaces and calculated the “transformational efficacy index” (TEI) for that trial’s representational transformation into the *Sample 1* subspace and its representational transformation into the *Sample 2* subspace. TEI was defined as the Euclidean distance between that trial’s representation in the subspace and the template representation, normalized by the distance between the two template representations in that subspace. (E.g., for trial *n*, the TEI for the *Sample 1* subspace would be the Euclidean distance between the trial representation projected into the *Sample 1* subspace and the *Sample 1* template (projected into the *Sample 1* subspace), divided by the distance between the *Sample 2* template projected into the *Sample 1* subspace and the *Sample 1* template (projected into the *Sample 1* subspace). A lower average TEI for a context-based transformation (e.g., the *Sample 1* transformation) corresponded to lower variability, for that subject, of that transformation, which we interpreted as higher “transformational efficacy.” For the fMRI data, we used TR 5-7 to define the *Delay 1.1* subspaces, and for the for the EEG data we used the entirety of *Delay 1.1*.

For priority-based transformations, we followed the same procedures, but used TR 9-11 to define the *Delay 1.2* subspaces and the entirety of *Delay 1.2* period for the EEG data, and labeled the data according to priority status (i.e., PMI and UMI).

If the efficacy of a context-based transformation is important for behavior, smaller TEIs should be associated with superior performance. To assess this in the fMRI data, for each subject we sorted responses, separately for *Recall 1* and for *Recall 2*, by median split of angular error, then calculated, for each response, the average TEI for each type of transformation (e.g., “what was the average TEI for the transformation to *Sample 1* for low-error vs. high error responses to *Recall 1*?”). Then we performed paired-samples *t*-tests between group-average high-error and low-error TEIs, separately for each subspace, each brain region, and each response (*Recall 1* and *Recall 2*). The analysis procedure was similar for the EEG data except that the comparison was between incorrect and correct responses.

To test how the TEI for UMI and PMI covary, we ran two-sided Spearman’s rank correlations between the two metrics across all trials for each subject and counted the number of subjects with correlations reaching the significance level of α = .05.

#### Within- and cross-label decoding of RNN and fMRI data

To assess where in the brain context and priority are represented, we carried out a series of decoding analyses applying the following logic. If a region represents context, any given stimulus item will be represented differently when it has the status of, for example, *Sample 1* versus when it has the status of *Sample 2*. If a decoder applied to data from this region can be successfully trained to classify stimulus identity when the data are labeled as *Sample 1* (successful “within-label” decoding), it should fail to decode stimulus identity when the data are relabeled as *Sample 2* (unsuccessful cross-label decoding). If a region that does not represent context, in contrast, any given item’s representational format will not differ as a function of its context status, and so a decoder that can be successfully trained on the data labeled as *Sample 1* should succeed at decoding stimulus identity when the data are relabeled as *Sample 2* (successful cross-label decoding). Before carrying out these analyses, we assumed the results with RNNs will have demonstrated that they can be trained to perform the DSR task. Therefore, it would necessarily be true that they represented both context and priority, and so applying these analyses to the RNN data would act as a sanity check for this logic. To be consistent with the fMRI dataset, RNN data were generated by testing the trained network on 324 trials of 9 possible orientations (counterbalanced across the identities of *Sample 1*, *Sample 2*, *Cue 1* and *Cue 2*, to be analogous to the Yu, Teng and Postle (2020) task), and subsequently extracting the RNN hidden layer activity. For the RNN data we decoded orientation and for the fMRI data we decoded location. (Decoding item location is generally more sensitive than decoding item orientation, and so demonstrations of failures of cross-label decoding of item location would provide stronger evidence for the encoding of the stimulus property of interest.)

For the RNN data and for the fMRI data from each ROI, we trained linear Support Vector Machine (SVM) multiclass classifiers to decode stimulus identity with a *k*-fold cross-validation procedure and a ‘one vs one’ coding design (see Supplementary Materials S1 for comparisons with results from other decoding methods). For context-based decoding, for each subject and at each timepoint, we trained a classifier with the data labeled as *Sample 1* then tested it on the data labeled as *Sample 1* (within-label decoding) and with the data labeled as *Sample 2* (cross-label decoding). We then repeated this process by training on *Sample 2*, and with fMRI data, for simplicity, we averaged the results to generate the overall accuracies for within-label decoding and cross-label decoding. For priority-based decoding, we used the same procedure except that the labels were *PMI* and *UMI*, instead of *Sample 1* and *Sample 2*, the *PMI/UMI* label reassigned at timestep 301 (for RNN) or TR 15 (for fMRI) to reflect identity of *Cue 2* (i.e., to account for the fact that priority status changed partway through “switch” trials). For the fMRI data, to evaluate the significance of decoding accuracy against chance level (1/9), we performed one-tailed one-sample *t*-tests against 1/9 on decoding accuracies across all subjects, and corrected for multiple comparisons using the false discovery rate (FDR) method.

## Results

### RNN

#### PCA visualization of hidden layer activity

PCA was carried out on the RNN hidden layer activity across all timepoints from 1000 withheld testing trials with *Sample 1* and *Sample 2* spanning the [0°, 180°) angular range and the resultant dimension-reduced activity projected into 3 subspaces that were spanned by PC1-PC2, PC2-PC3 and PC3-PC4, respectively, on a timepoint-to-timepoint basis. We trained three RNNs using the same training regime, used the PCA visualization of hidden layer activity from the first two for hypothesis generation and validation, and report results from the third network. The dynamical representational patterns observed in all 3 networks were highly consistent (See Supplementary Movie 1).

Upon the presentation of *Sample 1*, its representation formed a ring in the subspace spanned by the first 2 PCs, with relative distances between stimulus values preserved (as shown by the smooth color gradient of the ring; Figure 3A, top left panel), such that stimulus value can easily be read out from this subspace. Although there are also smooth color gradients in the other two subspaces, their geometry is more complex, making it less clear if they would support readout. The ring structure in the PC1-PC2 subspace was maintained across the ensuing ISI (see Figure 3A and Supplementary Movie 1). After the presentation of *Sample 2*, *Sample 2*’s identity was represented in the PC1-PC2 subspace, also in the form of a ring with a smooth color gradient (although the ring was somewhat “stretched out” relative to timestep 99; Figure 3B, bottom left panel). In parallel, information about *Sample 1* emerged in the subspace spanned by PC3 and PC4, in the shape of a ring with a smooth (albeit “stretched out”) color gradient (Figure 3B, top right panel). In effect, whereas *Sample 1* was represented in the PC1-PC2 subspace when it was the only item in WM, it was shunted to the PC3-PC4 subspace with the presentation of *Sample 2*, which replaced *Sample 1* in the PC1-PC2 subspace. Thus, prior to cuing, the RNN encoded the ordinal context of *Sample 1* and *Sample 2* by segregating them in orthogonal subspaces.

**Figure 3.**
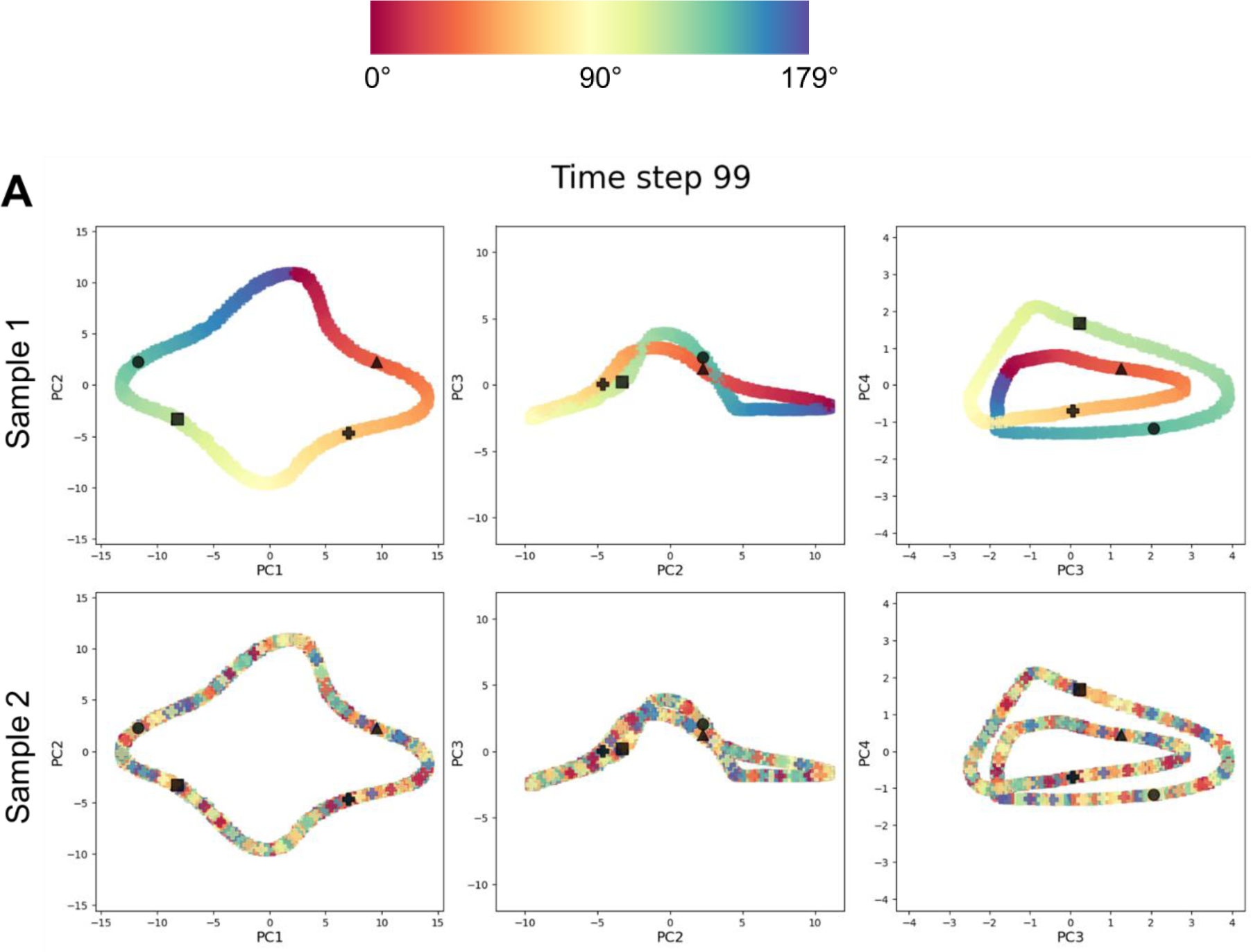
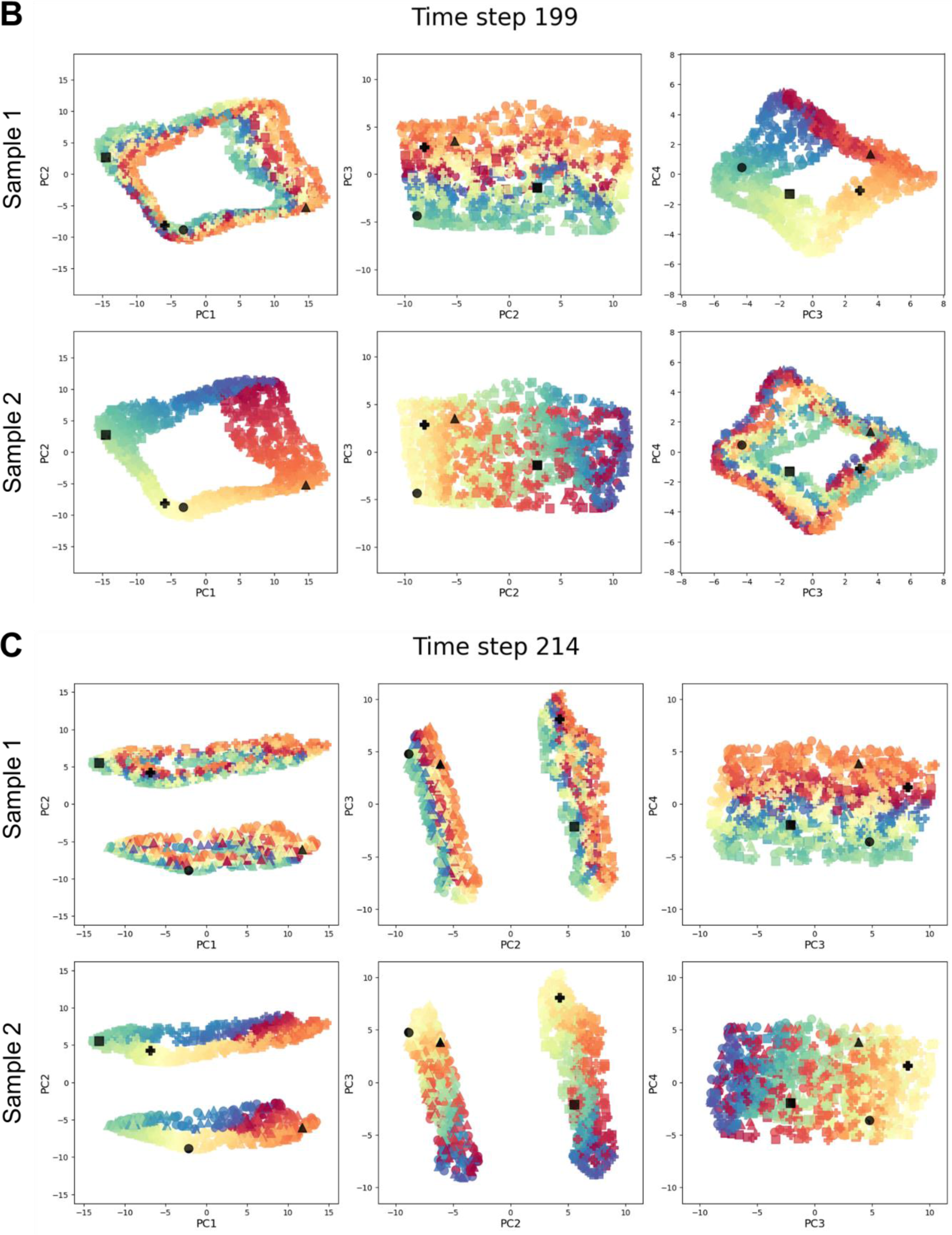
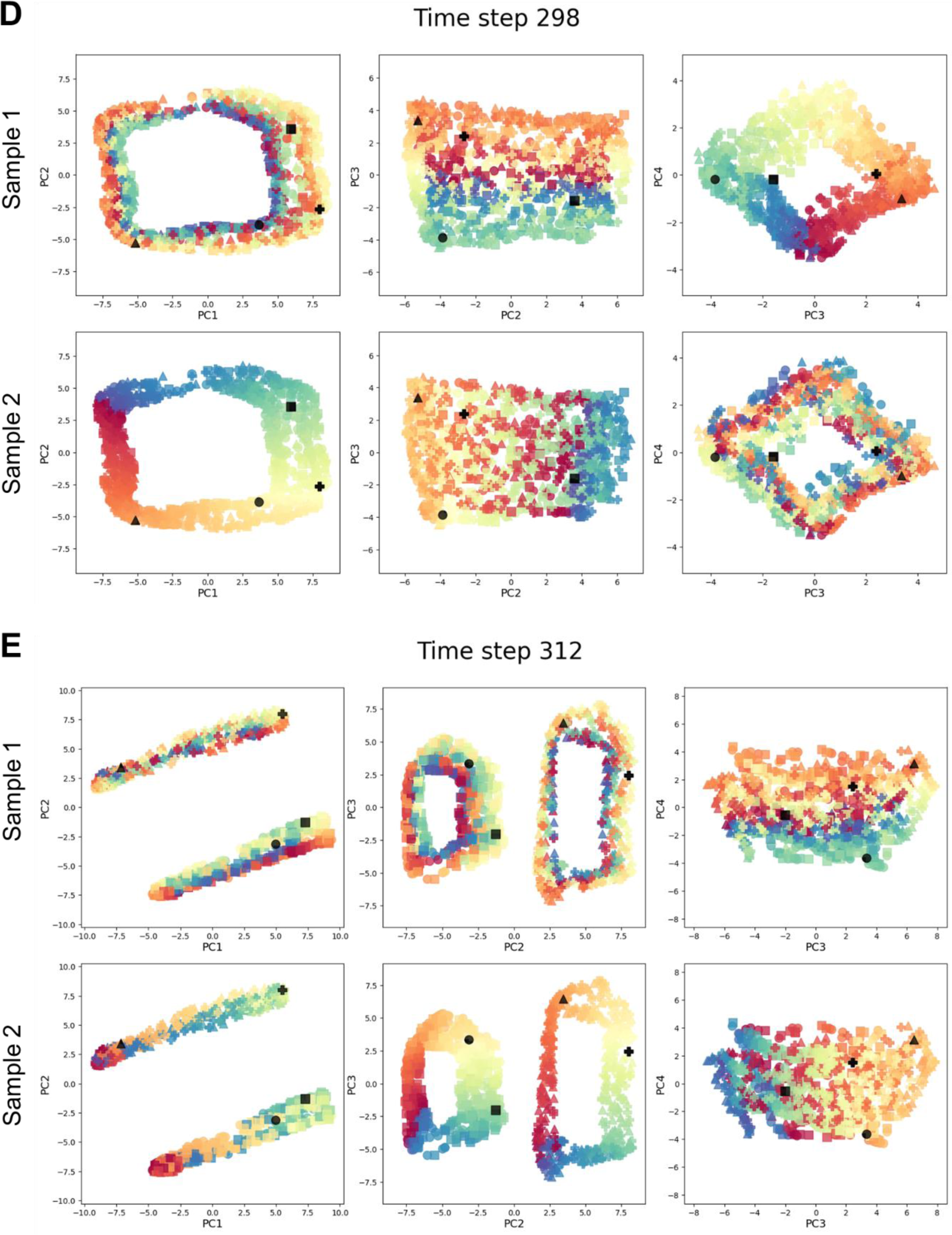
Visualization of representational dynamics embedded in RNN hidden layer at each of five representative timesteps across the DSR task. Each plot contains 1000 data points, one corresponding to each simulated trial, and the symbols indicating that trial’s cue configuration: *Cue 1* -> *Sample 1*, *Cue 2* -> *Sample 1*(●); *Cue 1* -> *Sample 1*, *Cue 2* -> *Sample 2* (▴); *Cue 1* -> *Sample 2*, *Cue 2* -> *Sample 1* (▪); *Cue 1* -> *Sample 2*, *Cue 2* -> *Sample 2* (+). In each plot, an example trial of each cue configuration is colored black for better visualization. For each of the five timesteps the same data are illustrated in six ways: the top row with the data labeled as *Sample 1* and the bottom row with the data labeled as *Sample 2*, and for each they are projected into three subspaces. A. After the presentation of *Sample 1* (Timestep 99). Note that because *Sample 2* has not yet been presented, the stimulus values are haphazard. B. After the presentation of *Sample 2* (Timestep 199). With both items in WM, but prior to cuing, *Sample 1* is now represented in the PC3-PC4 subspace and *Sample 2* in the PC1-PC2 subspace. C. During presentation of *Cue 1* and generation of *Response 1* (Timestep 214), illustrating a separation-by-priority status in the PC1-PC2 subspace. (A comparable priority-based separation was visible in the PC3-PC4 subspace earlier during this same epoch (not shown).) D. During the delay between *Cue 1* and *Cue 2* (Timestep 298). E. During presentation of *Cue 2* and generation of *Response 2* (Timestep 312), again illustrating a separation-by-priority status in the PC1-PC2 subspace but now based on *Cue 2*. (As with the *Cue 1* epoch, a comparable priority-based separation was visible in the PC3-PC4 subspace earlier during this *Cue 2* epoch (not shown).)

Upon the presentation of *Cue 1* (at timestep 201), the stimulus representations within each subspace separated into two clusters that were defined by priority status. For example, Figure 3C illustrates that in the PC1-PC2 subspace, at timestep 214, trials for which *Sample 1* was cued (denoted by triangle and circle symbols) separated from trials for which *Sample 2* was first cued (square and plus-sign symbols). Throughout the *Cue 1* epoch the axis along which this separation occurred rotated in multidimensional space over time. Thus, whereas timestep 214 was selected for Figure 3C because it clearly shows this separation-by-priority status in the PC1-PC2 subspace; the separation was visible in the PC3-PC4 earlier during this epoch, at timestep 207 (see Supplementary Movie 1). Thus, the RNN encoded priority status via separation within each subspace.

During the delay between *Cue 1* and *Cue 2* (timesteps 251-300), the prioritization clusters merged such that, prior to the presentation of *Cue 2*, information about *Sample 1* and *Sample 2* was again clearly observed in the PC3-PC4 subspace and in the PC1-PC2 subspace, respectively (Figure 3D). Finally, upon the presentation of *Cue 2*, the network representation once again separated into two priority-defined clusters, this time based on *Cue 2*’s identity, (i.e., trials for which *Sample 1* was cued (denoted by circle and square symbols) and trials for which *Sample 2* was cued (triangle and plus-sign symbols) separated into two clusters; Figure 3E). Thus, visualization of the representational of the RNN recurrent unit activities revealed that context and priority were represented via different transformational mechanisms, the former via the *segregation* of stimuli to orthogonal subspaces, and the latter via *separation* within each subspace.

#### Effective Dimensionality

During the processing of *Sample 1*, effective dimensionality (ED) initially rose to a value between 3 and 4 before declining to a value of ≈ 2 during the ensuing ISI (Figure 4). Upon the presentation of *Sample 2*, ED rose precipitously to a value close to 6 before declining steadily for the remainder of this epoch and the ensuing *Delay 1.1* to a value just below 3, which corresponds well to the encoding of a new stimulus and the segregation of subspaces to represent the ordinal context. The three remaining trial epochs were characterized by an initial increase of ED to a value of roughly 4 followed by a decline back to roughly 3. Particularly noteworthy in these results is the increase in ED following the offset of *Cue 1*. Note that because a similar increase in ED was not observed upon the offsets of the *Stimulus 1* or *Stimulus 2* epochs, this effect cannot be simply due to a transition from one epoch to the next. Rather, this effect closely resembled those time-locked to the onset of *Cue 1* and to the onset of *Cue 2*, events that each prompted the separation of stimuli into priority-defined clusters (Figure 3C and 3E). Therefore, it may be that the operation of removing from the network the encoding of no-longer-relevant information about priority status related to *Cue 1* -- corresponding to the merging of priority-defined clusters that was observed in the PCA visualization – is also an operation that entails a transient increase in ED.

**Figure 4.**
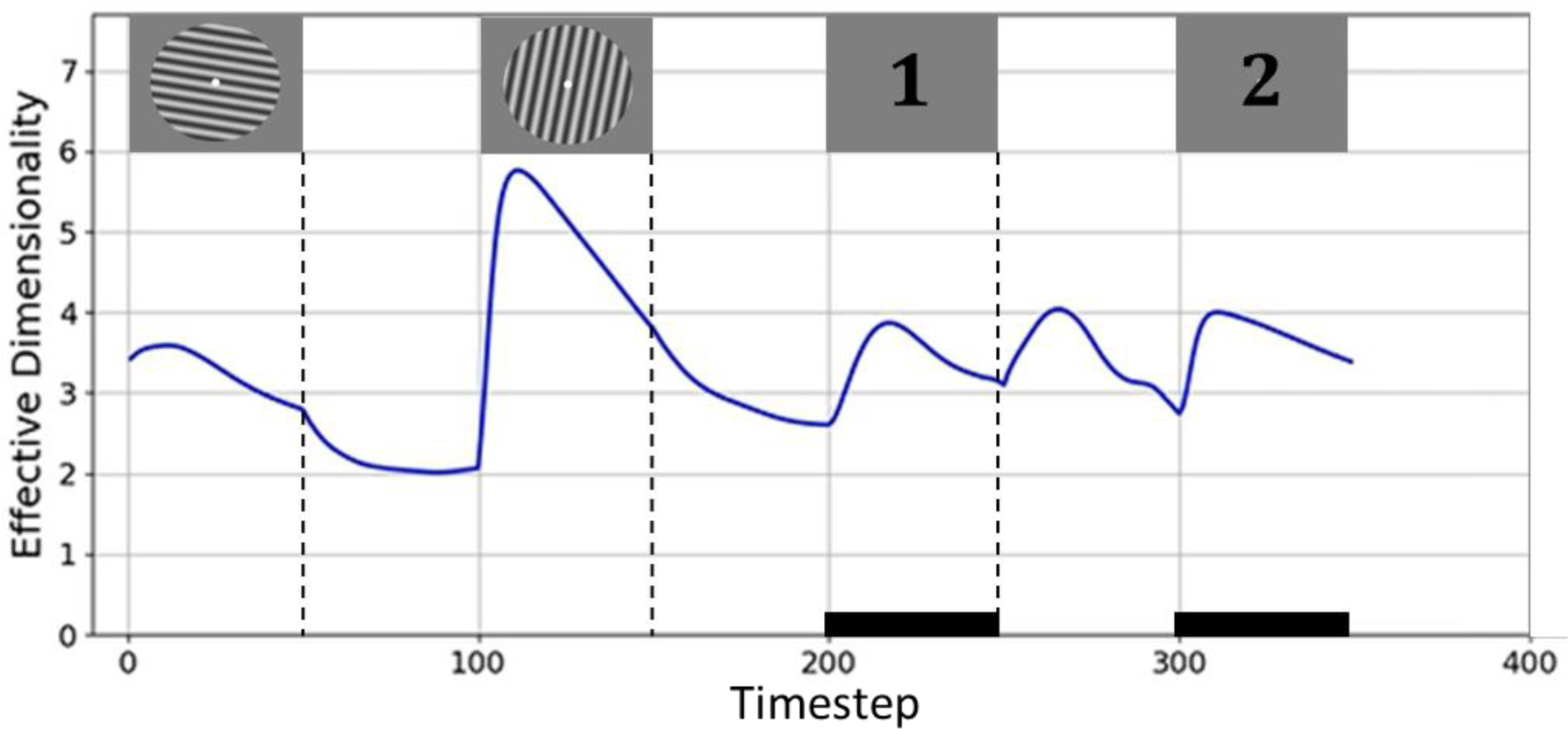
The time course of effective dimensionality (ED) of the RNN hidden layer stimulus representations. The rectangular images above the curve denote corresponding task events. The black rectangles along the *x*-axis indicate time periods when a response was being made.

### Interim Discussion

We trained RNNs to perform the DSR task and applied dimensionality reduction to the internal representations of the network. Visualization of the representational dynamics yielded several important insights. First, prior to the first prioritization cue, information corresponding to the two sample stimuli was represented in orthogonal subspaces (Panichello & Buschman, 2021). This may serve not only to individuate the two, but to encode the distinct ordinal context that the network needed to correctly interpret the cues. Second, priority status was represented by separating stimulus representations to distinct regions within each subspace, according to the cued “context”. The first observation is important because it emphasizes the importance of encoding trial-specific context in WM, and previous empirical studies of prioritization have largely overlooked this operation. The second observation is important because it indicates that the representation of priority status may be implemented in a different way, via separation within a subspace, than is ordinal context, via segregation of stimulus information to distinct subspaces. This difference is all the more interesting when one considers that to the RNN, ordinal position and priority may be just two dimensions of task context that play out on different time scales during a single trial. In this variant of DSR, one dimension of an item’s context is the order in which it was presented. This can be considered the “first-order” context because it uniquely individuates an item for the duration of a trial, and it does not change for the duration of the trial. (It is to first-order context that Oberauer and Lin (2017) refer when they state that the binding of context to a stimulus is fundamental to that stimulus being in the state of being “in WM”.) A second dimension of context is priority status, and this differs from first-order context because its status for an item varies within the trial between “not applicable,” “prioritized,” and “unprioritized” (indicated by values of 0, 1, and -1, respectively, being input by the cue unit). Thus, priority serves as a “second-order” level of context, one that indicates an item’s in-the-moment status with respect to the rules of the task, and that cannot be interpreted in the absence of information about first-order context. These considerations highlight that to fully understand the flexible control of WM we need to understand how first-order context is coded in the brain and how it interfaces with higher-order context to guide thought and action.

Recent empirical studies that have manipulated demands on first-order context in WM have implicated regions of frontal cortex and the intraparietal sulcus (IPS) (for ordinal context, see Fulvio et al., 2023; Gosseries et al., 2018; for location context, see Cai et al., 2020; Fulvio et al., 2023). In Yu, Teng & Postle (2020), a study that also manipulated priority, the location context of differently prioritized orientation stimuli was found to be preferentially coded in IPS, and not early visual cortex, even though location information was not directly tested by the task. More recently, Teng and Postle (Stage 1-accepted registered report) used the same stimuli and procedure, but flipped the roles of context and content, making orientation the first-order context used to cue memory of an item’s location. “Context load” was manipulated via the similarity of orientation of the two sample stimuli, and individual differences in context-load sensitivity of activity in IPS (but not early visual cortex) predicted behavioral sensitivity to this factor. Generalizing across these studies suggests that first-order context in WM may be represented more prominently in areas associated with cognitive control than in areas associated with stimulus representation. The same may not be true for second-order context, because prioritization effects are prominent in early visual cortex (Yu, Teng & Postle, 2020; Teng & Postle, Stage 1-accepted registered report).

These considerations, prompted by the results from the RNNs, highlighted for us the importance of understanding the encoding of first-order context in WM, and of understanding similarities and differences of neural and behavioral correlates of first-order versus second-order context. What follows are initial attempts to do so, via reanalyses of an extant fMRI and an extant EEG dataset from two previous studies of the DSR task.

### Analyses of fMRI and EEG data

The fMRI study used a DSR procedure that was most closely matched to that used with the RNN, including the fact that it used stimulus order as first-order context. The fMRI data would also allow for assessment of possible regional differences in the representation of the two types of context. The task used in the EEG study used location as the dimension of first-order context, and so would allow an assessment of generalization of what has been observed for ordinal context (with the RNN and fMRI) to location context. (For ease of exposition, in the results that follow we will refer to first-order context as “context” and second-order context as “priority,” because priority is the only dimension of second-order context that is relevant in the DSR task.)

### Transformational efficacy

One way to compare the neural representation of context versus priority is to assess their influence on behavior. To do this, we took an individual differences approach, using the variability of trial-by-trial encoding of context and of priority as proxies of the efficacy with which these operations were carried out. (I.e., a subject for whom context-based or priority-based transformations were more variable from trial-to-trial might be expected to perform worse on the task.)

For context, results failed to show evidence that behavior was sensitive to transformational efficacy. For the fMRI data (ordinal context), transformational efficacy indices (TEI) did not differ for low-vs. high-error trials, for *Recall 1* or *Recall 2*, in any of the 3 ROIs (early visual cortex, IPS 0-5, FEF; all *t*(12) < 1.74, *n.s.*). For the EEG data (location context), TEI did not differ for correct vs. incorrect trials (*t*(11) < 1.37, *n.s.*).

For priority, there was considerable evidence that behavior was sensitive to transformational efficacy. For the fMRI data, in early visual cortex, TEI was lower for low-error than high-error trials for the UMI subspace for *Recall 1* (*t*(12) = 1.81, *p* < .05) and for the PMI subspace for *Recall 2* (*t*(12) = 2.06, *p* < .05). For IPS0-5, TEI for the PMI subspace was lower for low-error than high-error trials for *Recall 1* (*t*(12) = 2.04, *p* < .05), and was lower for low-error than high-error trials for both UMI (*t*(12) = 3.04, *p* < .01) and PMI (*t*(12) = 3.00, *p* < .01) subspaces for *Recall 2*. All other comparisons, including all for FEF, failed to achieve significance (all *t*(12) < 1.57, *n.s.*). For the EEG data, TEI was lower for correct trials than incorrect trials for *Recall 1* in both UMI (*t*(11) = 2.17, *p* < .05) and PMI (*t*(11) = 4.28, *p* < .001) subspaces.

The TEI also offered a metric with which to begin exploring whether the transformation to PMI and the transformation to UMI may share a common component that acts on the two simultaneously (c.f., Panichello & Buschman, 2021). Specifically, we correlated trial-by-trial TEI for the PMI with trial-by-trial TEI for the UMI (two-sided), reasoning that evidence of correlation would be expected if the two do share an underlying mechanism. For the fMRI data, in early visual cortex, this correlation was significant at *p* < .05 for 12 out of 13 subjects, in IPS 0-5 for 11 subjects, and in FEF for 10 subjects. For the EEG data, TEIs for PMI and UMI were significantly correlated for 10 out of 12 subjects. All correlations were positive.

Within- and cross-label decoding of RNN and fMRI data

#### RNN data

To assess the representation of both context and priority in the RNN, we performed within- and cross-label decoding analyses on the RNN recurrent unit activities across all 350 timesteps from 324 novel, counterbalanced trials of 9 different orientations using a linear SVM classifier (Figure 6).

**Figure 5.**
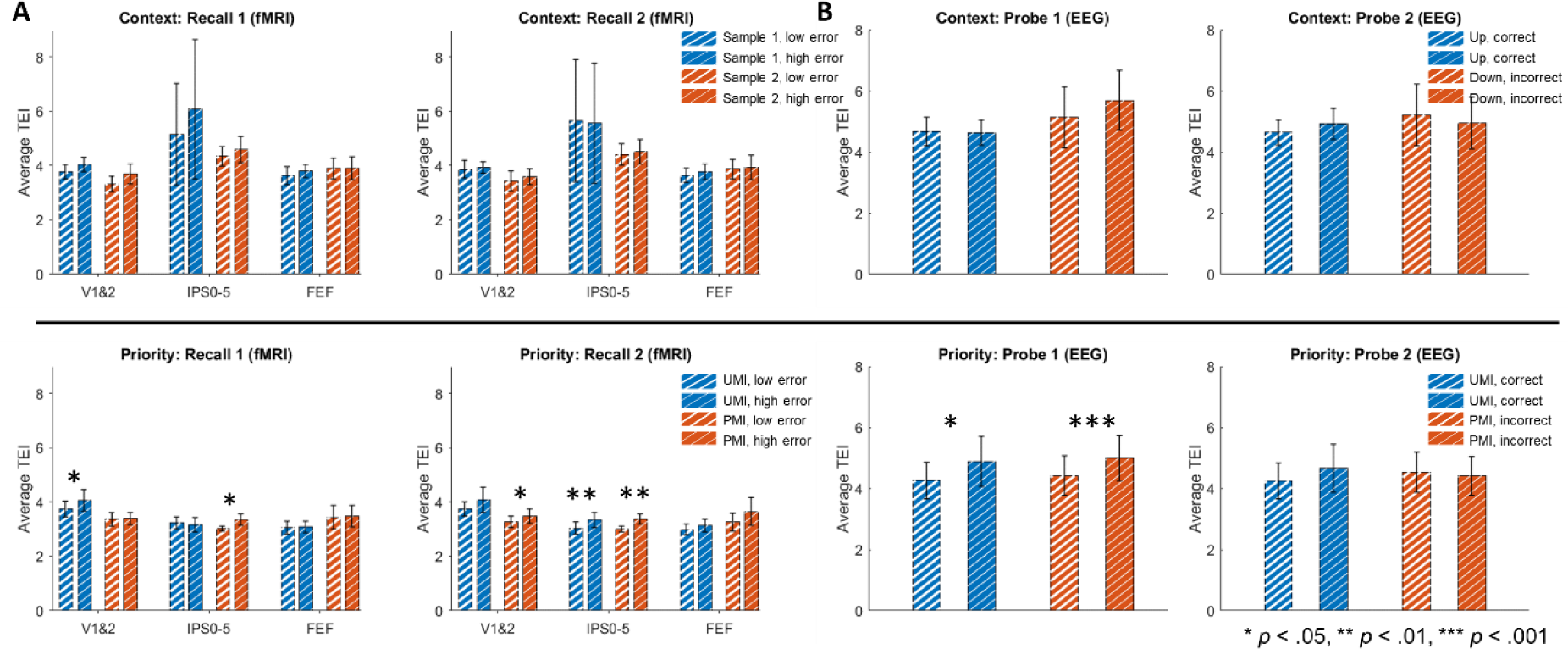
Transformational efficacy analysis results on fMRI (Yu, Teng & Postle, 2020) and EEG (Fulvio & Postle, 2020) data. (A) Comparisons between average TEI for high-error and low-error trials across subjects from the fMRI dataset. (B) Comparisons between average TEI for incorrect and correct trials across subjects from the EEG dataset. Top row: priority-based decoding; bottom row: context-based decoding. The subspace from which the TEI is calculated is indicated in the legends. Asterisks above bars of the same color indicate the significance level of the paired-sample *t* tests comparing the average TEI between each two groups.

**Figure 6.**
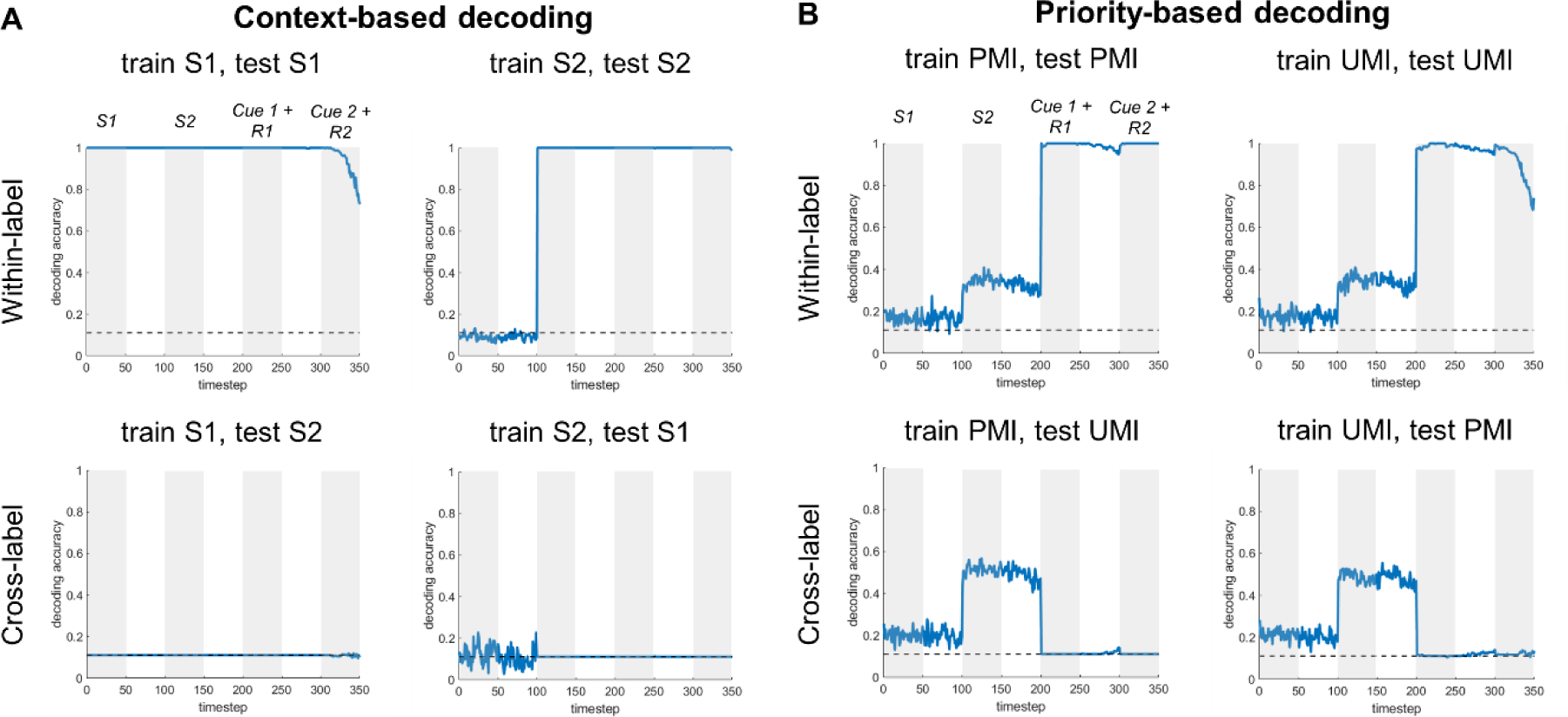
Within- and cross-label decoding of RNN data across the DSR trial. (A) Context-based decoding. Classifiers were trained on *Sample 1/2*, then tested on *Sample 1/2* (within-label), or tested on *Sample 2/1* (cross-label). (B) Priority-based decoding. Classifiers were trained on PMI/UMI, and tested on PMI/UMI (within-label) or tested on UMI/PMI (cross-label). S1: *Sample 1*, S2: *Sample 2*; R1: *Response 1*; R2: *Response 2*.

For context-based decoding we obtained close to perfect decoding accuracy when training and testing on the labels of the same sample throughout the task (note that for train S2, test S2 decoder performance was at chance prior to timestep 101, due to the absence of information about *Sample 2* at those time steps). For cross-label decoding, however, accuracy was at chance level for the duration of the trial. For priority-based decoding, within-label decoding accuracy for both PMI and UMI was close to chance level prior to *Cue 1*. With the onset of *Cue 1*, for both PMI and UMI, decoder performance rose to close-to-perfect for the remainder of the trial. For cross-label decoding, whereas decoding accuracy for both PMI and UMI was above chance level prior to *Cue 1*, for both it dropped to chance level with the onset of *Cue 1*, and remained there for the remainder of the trial. Both of these sets of results validated the reasoning that a system that represents context and priority would not support cross-label decoding for either factor.

#### fMRI data

We investigated the anatomical distribution of the representation of context and priority during the DSR task by carrying out a series of multiclass decoding analyses on the fMRI dataset. In general (and unlike for the RNNs) decoder performance was far from ceiling, and tended to be superior for time points corresponding to trial epochs when stimuli were on the screen. Importantly, however, we were generally able to decode the stimulus identity across the whole time course with above-chance accuracy in every ROI, especially in the time period between *Cue 1* and *Cue 2*, where one stimulus is prioritized over the other in working memory (within-label rows of Figure 7). (The one exception was in IPS4 with context-based decoding; the reason for this is unclear.)

**Figure 7.**
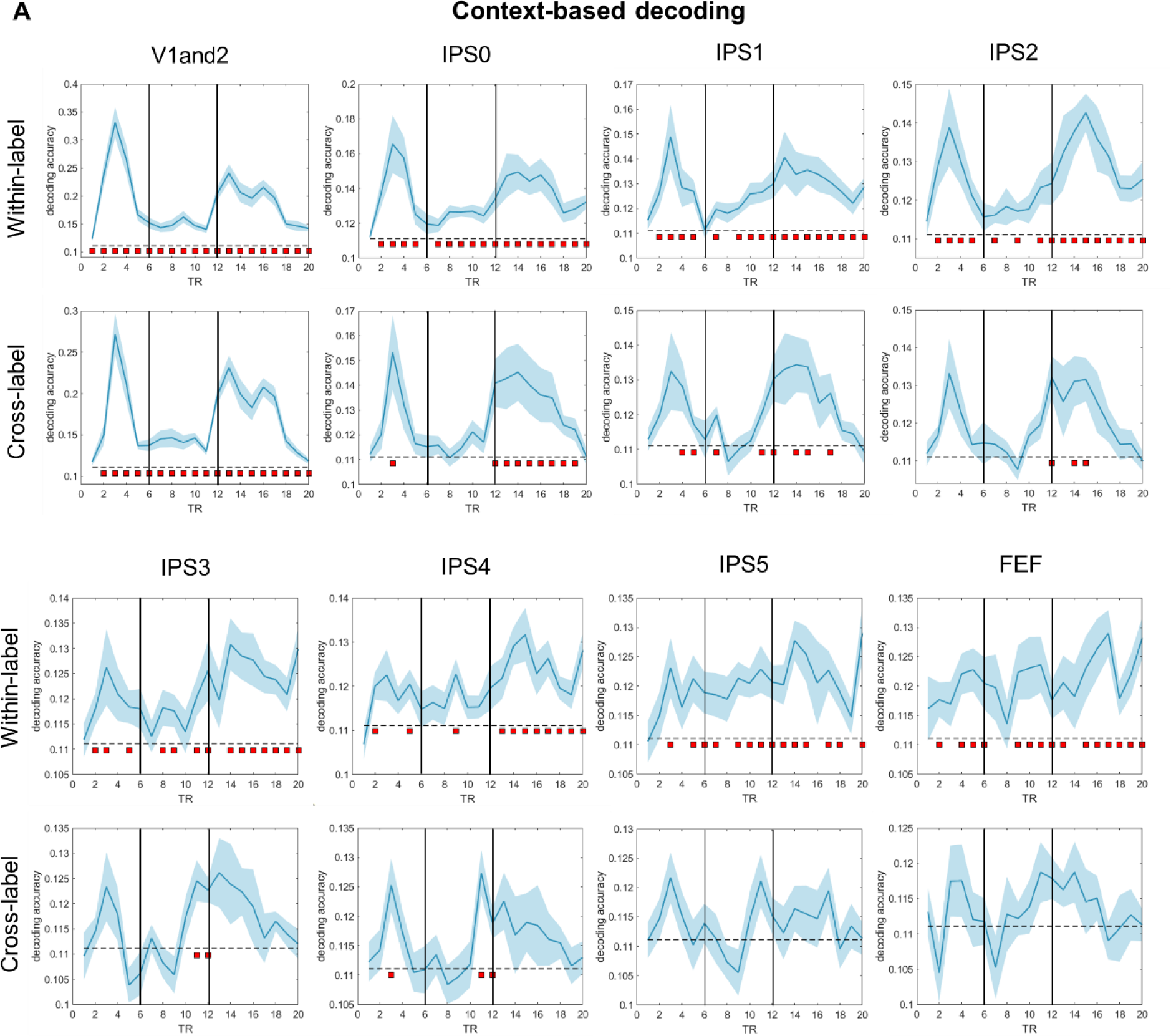
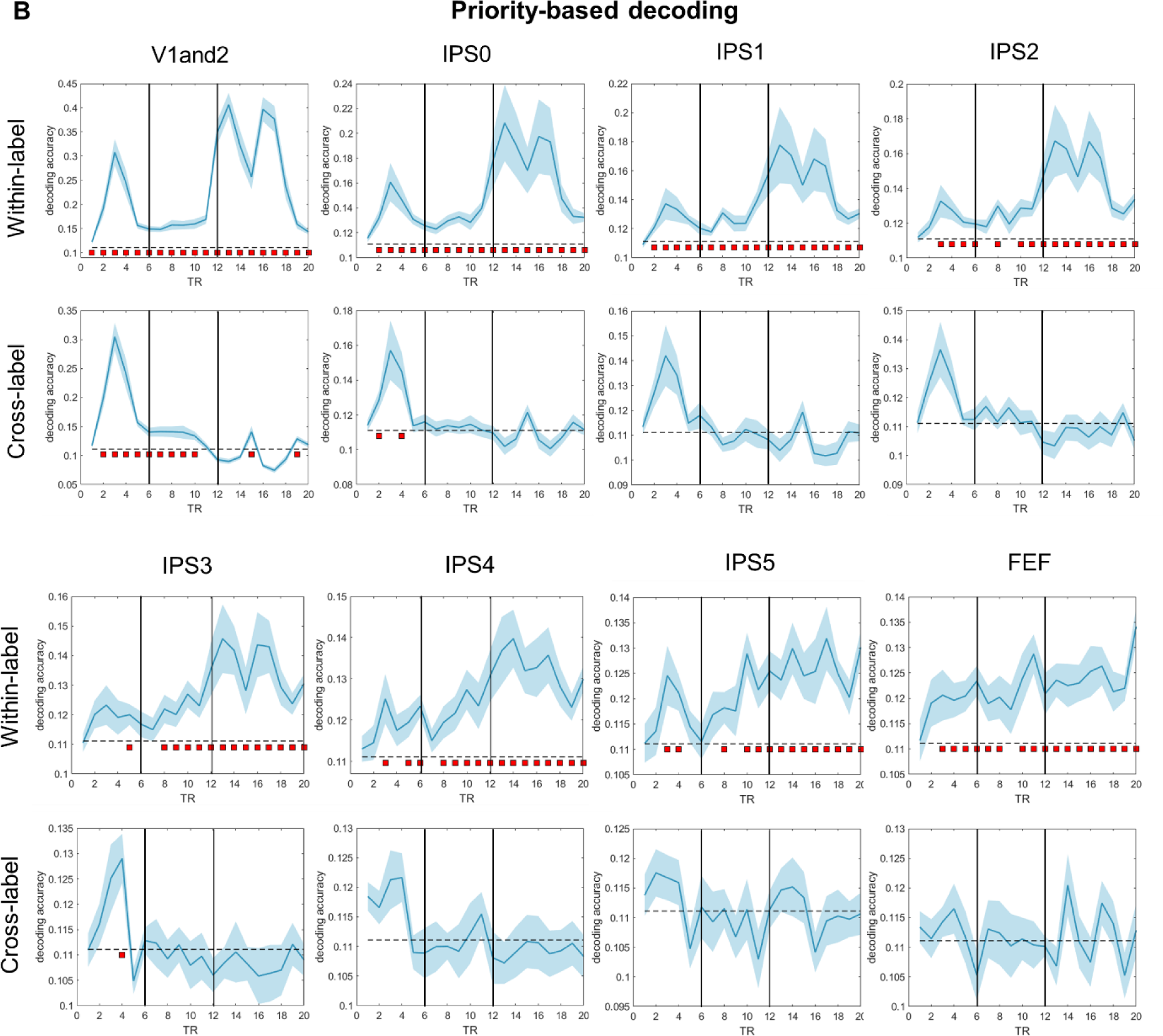
Within- and cross-label decoding analyses from the fMRI dataset. (A) Context-based decoding. (B) Priority-based decoding. In each graph, the two vertical solid black lines indicate *Cue 1* and *Cue 2*, respectively. The blue shading around each curve shows standard error of the mean. The horizontal dashed line indicates the chance-level decoding accuracy of 0.11. Red squares below the dashed line indicate time points with significant above-chance decoding accuracy (*p* < .05, FDR-corrected across all time points). Note that the range of the *y*-axis varies from graph to graph.

For context, cross-label decoding revealed a marked posterior-to-anterior gradient: it was successful for the entirety of the trial in V1-V2 (indicating that changing the context does not affect the decoding accuracy, and hence, context information is likely ignored); successful for *Cue 2* and *Delay 2* epochs for IPS0 and for a smaller number of timepoints for IPS1 and IPS2; successful only for late *Delay 1.2* for IPS3 and IPS4, and entirely at chance for IPS5 and FEF (Figure 7A). This indicates that context was not represented at the earliest stations of the visual system and become progressively more prominent at progressively higher levels of the dorsal stream.

For priority, cross-label decoding for V1-V2 was successful for the beginning of the trial through late *Delay 1.2*, after which it dropped to baseline. For the remainder of the ROIs cross-label decoding was at baseline for the entirety of the trial. This suggests that priority is represented in every ROI that we investigated, albeit taking longer to manifest in V1-V2 (Figure 7B).

## Discussion

In this study we initially set out to investigate the mechanisms underlying prioritization on a task in which changes of priority were not predictable – the double serial retrocuing (DSR) task – via visualization of representational dynamics of an RNN trained to perform the task. Unexpectedly, results from the RNN called to our attention the importance of also understanding the representation of an additional dimension of trial-specific information, the first-order context that uniquely individuates each item during the trial. Across model training, validation and testing, we saw that the encoding of first-order context was accomplished via the segregation into orthogonal subspaces of the representation of the first and second item to be presented. Unlike first-order context, higher-order context can change within a trial, a property that is often manipulated with instructional cues. In the DSR, priority status is the second-order context, and it is specified, then removed, then specified a second time, during the course of each trial. The encoding of priority was accomplished via separation within each context-encoding subspace of prioritized from unprioritized items. Thus, the RNN indicated that first- and second-order contexts are encoded via distinct mechanisms, *segregation* to orthogonal subspaces versus *separation* within a subspace, respectively. Furthermore, an effective dimensionality analysis suggested that the operation of resetting second-order context (as happens during the ISI separating *Cue 1* and *Cue 2* in the RNN version of the task) may make computational demands that are comparable (in terms of requiring additional dimensions) to those needed to establish it.

Consistent with the distinct dynamics observed with RNNs, reanalyses of an fMRI and an EEG dataset established that the processing of first- and second-order contexts has distinct behavioral and neural profiles for humans performing the DSR. To assess relations to behavior, dimensionality reduction was applied to the neural data and transformational efficacy indices (TEI) derived for each subject for each of the two levels of first-order context and for each the two levels of second-order context. Correlations with behavior failed to show any evidence that performance is sensitive to variation in TEI for first-order context, whether first-or-second-to-be-presented (fMRI study) or top-or-bottom-location-of-presentation (EEG study). For priority (i.e., second-order context), in contrast, there was considerable evidence that larger TEIs (indicating higher trial-to-trial variability) corresponded to poorer performance. In the fMRI dataset, the anatomical distribution of the representation of order and priority also differed, with the former absent from early visual cortex and becoming progressively more robust in more rostral ROIs, whereas the latter was evident in every ROI that we investigated. Thus, our results suggest that not only are representational transformations corresponding to first-order versus second-order context implemented via different mechanisms, they also differ according to their influence on behavior and to their distribution in the brain.

These results share some similarities and some differences with a recent study that recorded neuronal activity from several brain areas of nonhuman primates (NHP) performing a single-retrocue working memory task (Panichello & Buschman, 2021). In that study, dimensionality reduction revealed that, prior to the retrocue, the two stimuli were represented in orthogonal subspaces that corresponded to the location at which each had been presented (i.e., first-order context). Upon cuing, stimulus information transformed into different “post-selection” subspaces that retained first-order context and now also represented selection status (selected/non-selected; i.e. second-order context). Notably, the representations of “selected upper” and “selected lower” items were no longer orthogonal. The degree of cue-triggered representational transformation was highest in dorsolateral PFC and progressively weaker in more posterior regions, weakest in extrastriate visual area V4. One similarity of those results to those reported here is the encoding of first-order context into orthogonal subspaces. A notable difference between the two is the nature of the post-cue transformations.

In the DSR, the representational transformation of one item into a PMI and the other into a UMI are prompted by the same cue, a design feature that allows for direct comparison of the two processes. For the majority of subjects in the EEG study, and in the majority of ROIs in the majority of subjects in the fMRI study, trial-by-trial variation in the TEIs for the transformation to PMI and for the transformation to UMI were correlated, a result consistent with the idea that a common factor underlies both. There are at least two possible accounts for this pattern of results that will require future research to adjudicate. One is a parallel mechanism whereby a single signal is “split” so as to trigger the simultaneous output gating of one item into the PMI state and of the other item into the UMI state. A second is a serial process akin to biased competition (c.f., Desimone & Duncan, 1995) whereby a control signal first selects the cued item, and a consequence of this item’s transformation to PMI is that it “pushes” the other item into the UMI state. Importantly, the correlation of TEIs reported here rules out what had been a third possibility, which was a “passive” account of the transformation to UMI whereby the withdrawal of attention would allow the relaxation of the representation into a default state such that the relaxation process would not be influenced by the active PMI transformation. Along with the application of second-order context that is prompted by the prioritization cue, the time course of effective dimensionality of the RNN suggests that the resetting of second-order context partway through the trial may be a process that requires as much active control as does its initial application.

## Acknowledgement

We thank Dr. Yuri Saalmann and Jiangang Shan for their critical feedback. Simulations were partially performed using the CloudLab computing facilities (Duplyakin et al., 2019).

