## Supplementary Figure 1 for "Representing context and priority in working memory"

### Supplementary Materials

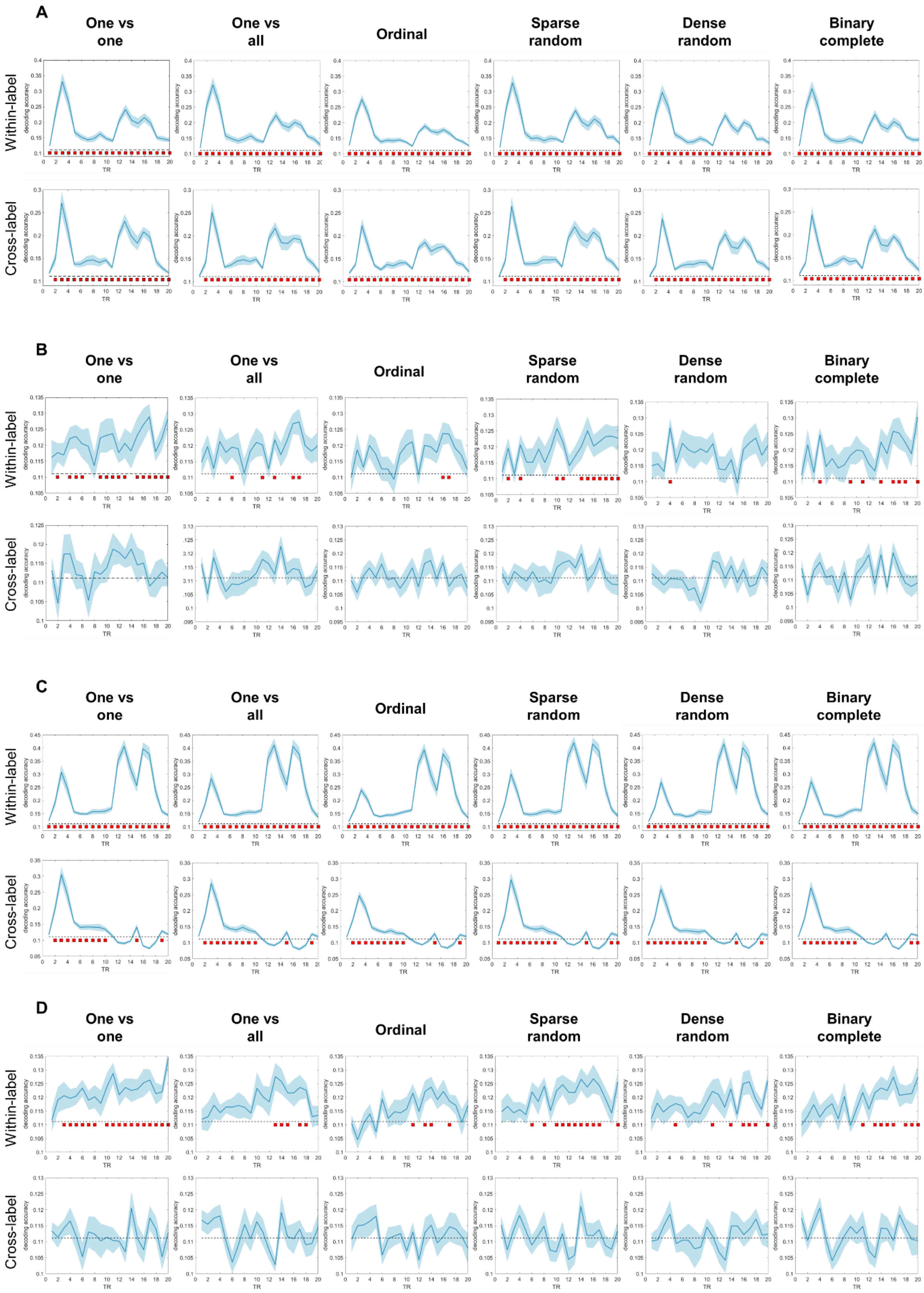

**Figure S1.** Comparisons of within- and cross-label decoding from the fMRI dataset across various SVM coding designs in MATLAB. (A) Context-based decoding for V1-2. (B) Context-based decoding for FEF. (C) Priority-based decoding for V1-2. (D) Priority-based decoding for FEF. In each graph, the blue shading around each curve shows standard error of the mean. The horizontal dashed line indicates the chance-level decoding accuracy of 0.11. Red squares below the dashed line indicate time points with significant above-chance decoding accuracy ($p < .05$, FDR-corrected across all time points).
